# Cytological abnormalities and pollen abortion in interspecific hybrids of *Nicotiana*

**DOI:** 10.1101/2019.12.12.873968

**Authors:** Jugou Liao, Zhiyun Chen, Zihui Pan, Yongzhi Niu, Wenlong Suo, Xuemei Wei, Yunye Zhen, Litang Lv, Wenguang Ma, Suiyun Chen

**Author notes:** Corresponding to: Suiyun Chen.

## Abstract

Interspecific cross breeding introduces superior agronomic traits into cultivated species; however, problematic pollen sterility occurs in the hybrids. Our previous study obtained interspecific hybrids from the cross between a cytoplasmic male sterility line of *Nicotiana tabacum* and *Nicotiana alata*, and some of the hybrids were pollen sterile. Here, we conducted an in-depth cellular study to understand the cytological mechanism of pollen abortion in these hybrids (F1-D sterile) compared with pollen development in the fertile hybrids (F1-S sterile) from the same cross. The ultrastructure observation showed that the membrane of microspore in F1-D sterile hybrid was deficient at all represented developmental stages. Chromosome behavior during meiosis was studied by carbol fuchsin staining, which indicated that cytomixis, chromosome leakage and asymmetric callose wall deposition occurred with high frequency in the microsporocyte of F1-D sterile. The results of the ultrastructure and 6-diamidino-2-phenylindole (DAPI) analyses also showed that the cytoplasm and nucleus were unstable and extruded from F1-D sterile microspore during the developmental process, leading to mature pollen grains that were vacuous and collapsed in the aperture region. In addition, delayed tapetum degradation was also detected in the anther of F1-D sterile, and might be associated with irregular sporopollenin deposition in the aperture region of F1-D sterile pollen. Genetic unbalance and cytomembrane deficiency might both be responsible for the instability of the chromosome, nuclear and cytoplasm, and resulted in pollen abortion in F1-D sterile hybrids, and irregular tapetum degradation might also be related with pollen sterility.

## Introduction

Hybrid sterility is a widespread postzygotic reproductive barrier between species or subspecies. This effect provides the initial force of genetic differentiation and speciation, and it represents a major barrier to the effective use of inter-subspecific genetic resources in many crops, including tobacco [1-3]. Pollen sterility is one of the most important reasons for interspecific hybrid sterility, along with embryo sac sterility[4, 5], and severely prevents the utilization of genetic diversity in hybrids; therefore, it is of great significance to understand the genetic or cytological mechanism of pollen abortion in hybrids.

The anther locule is where the pollen grain develops, and the developing anther consists of microsporocytes in the center, surrounded by the tapetum and the anther wall [6, 7]. The anther wall of *Nicotiana* belongs to the basic type and is composed of the endothecium, two middle layers and the tapetum [8, 9]. The tapetum, which is in direct contact with the microspores, serves as a nutritive and secretory tissue to supply the nutrients and other components required for pollen wall development, and it regulates the development of microspores during pollen mother cell (PMC) meiosis and the subsequent microspore maturation [9-11]. Pollen development in most of the species includes the following key stages: meiosis, tetrad and primexine development, microspore release, sporopollenin deposition, mitosis, accumulation of starch or lipids, addition of pollenkitt or tryphine to the exine, and dehydration of the cytoplasm [12-14]. Before the tetrad stage, the microsporocyte is encased in the callose wall, which is synthesized by the meiocyte and deposited between the plasmalemma and the original cellulosic wall during prophase I of meiosis [7, 15]. This callose layer is vital for preventing cell cohesion and fusion and useful as a template for the formation of species-specific exine-sculpting patterns; then, upon its dissolution, the free microspores were released [7, 16-20].

Male gametophyte development is prone to influence by the external environment and by inherent genetic and physiological conditions; therefore, the coordination between the anther sporophytic tissue and the microspore is crucial for pollen development and fertility. Pollen abortion may occur at various stages of pollen development. Primarily, chromosome behavior during meiosis must be under strict genetic regulation to facilitate the correct segregation of chromosomes into daughter cells, and abnormal meiosis inevitably induces pollen abortion [21, 22]. Scott reported that the absence of a callose wall is not necessarily fatal, and indeed, some species lack callose; however, irregular callose accumulation and dissolution are associated with male sterility in taxa where a callose wall is normally present [6, 21, 23]. Moreover, abnormality in the cell membrane might interfere with the movement of substances in and out of cells and with the development of the pollen wall structure [24]. Cell membrane expansion at the unicellular microspore stage is vital for pollen cell enlargement, polarization, mitosis, and finally pollen viability [1]. In addition, the timing of tapetum formation and degradation is critical for microspore development and pollen viability; premature or delayed tapetum degeneration causes male sterility [25, 26]. Aberrant pollen wall formation might also lead to pollen abortion. For example, impaired primexine formation often results in abnormal exine and aborted pollen [11, 27, 28].

Cytoplasmic male sterility (CMS) lines are a convenient and efficient tool for plant cross breeding, and the nuclear fertility restoration gene is necessary for fertility restoration in offspring obtained using a CMS plant as the maternal parent [29-32]. In a previous study, the CMS line of *Nicotiana tabacum* L. cv. (*gla.*) S ‘K326’ (with degenerated stamens, n=24) was hybridized with *Nicotiana alata*(n=9), which overcame the pre-fertilization cross-incompatibility encountered when using the corresponding male fertile line as the maternal parent, and F1 offspring with different morphology and fertility were obtained. Offspring named F1-S was morphologically similar to the maternal parent, with well-developed stamens and fertile pollen grains. However, hybrids F1-D was morphologically different from the female parent, with the stamens (anthers and filaments) developed, but the pollen grains were completely sterile [33]. We deduced that a fertility restorer from the paternal parent (*N. alata*) was responsible for male fertility restoration in F1-S fertile, and this hypothesis is undergoing further validation. However, another interesting question is why the F1-D sterile hybrids showed aborted pollen. To reveal the cause of pollen abortion in F1-D sterile, we studied the cytological mechanism of pollen abortion through analysis of pollen viability and fertility, microsporogenesis, callose wall deposition, cell ultrastructure, cytomembrane integrity, cytoplasm and nucleus stability, and tapetum development and degradation, which were compared with that in the F1-S fertile hybrids.

## Materials and Methods

The CMS line *Nicotiana tabacum* L. cv. (*gla.*) S ‘K326’ (n=24) was hybridized with *N. alata* (PI 42334, n=9) in June 2015. The materials were all provided by the Yunnan Academy of Tobacco Agricultural Sciences (Nanxiang Road 14, Yuxi, Yunnan, China). Offspring seeds were obtained in July and then germinated in January 2016 and grown for the next 5 months in the greenhouse of the Southern Tobacco Breeding Center, Yuxi, Yunnan. The plants were manually watered once a week and fertilized with water-soluble fertilizer (N: P: K 20:20:20+0.5% microelements) (Demei, Sichuan, China) once a month. Pollen grains from hybrids morphologically different from the maternal parent (F1-D-2 sterile) were utilized to explore the cytological mechanism of male sterility, and pollen from F1-S fertile (F1-S-2) was also studied as a control. The genome size of these two hybrids was previously reported [33]. Flowers were categorized into nine developmental stages. Stages 1 and 9 represented the smallest flower bud and the blooming flower, respectively. Information with images of each stage was available in Fig. 1.

**Fig. 1.**
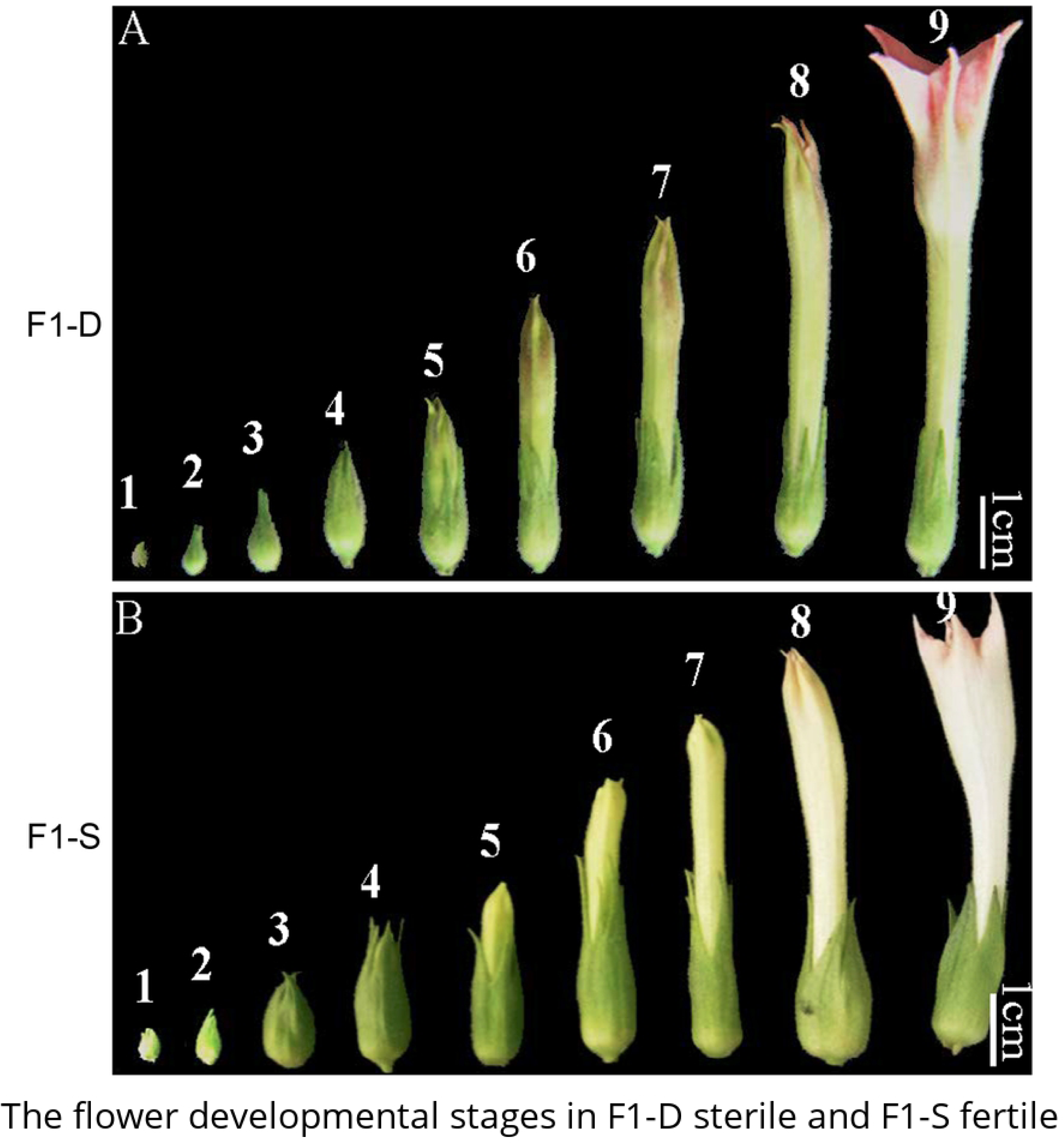
The flower developmental stages in F1-D sterile and F1-S fertile hybrid. The flower stage was corresponding to the development stages of PMC, microspore or pollen grains.

### Pollen viability and germination capacity estimated by FDA-PI and germination

Fluorescein diacetate (FDA) (Solarbio, Beijing, China) shows cell viability by detecting the presence of active esterases in the cytoplasm. Propidium iodide (PI) (Solarbio) could only permeate the membrane of dead cells, embed into double-stranded DNA in the nucleus and exhibit red fluorescence. FDA (10 mg) was dissolved in 20 μL of acetone and diluted to 5 ml with laboratory-pure water, and then PI was added to a final concentration of 1 mg mL^-1^. Microspores or pollen grains were collected from anthers at different developmental stages and stained with the above FDA-PI solution for 5 min. Signals were detected with a fluorescence microscope (Olympus BX 51, Olympus, Japan) by monitoring emission at 520 nm with an excitation wavelength of 485 nm. Only cells exhibiting bright green fluorescence were considered to be viable [23, 25, 34].

A basic in vitro germination medium (containing 10% sucrose, 0.01% boric acid and 0.01% Ca^2+^) was used to culture pollen at 28 °C for 3 h. A pollen grain was considered as germination when the pollen tube length equaled or exceeded the pollen grain diameter. Pollen germination capacity was also assayed by pollination. Pollen grains from the sterile and fertile hybrids were collected and deposited on the stigmas of the maternal parent line. Pistils were picked at 24 h after pollination and observed with fluorescence microscopy after aniline blue staining to detect pollen germination [35].

### Meiosis and callose wall observed by carbol fuchsin staining

Microsporocytes or microspores were collected from flower buds at stages 1 to stage 3, and anthers from each bud were placed on a glass slide. The microspores were released with the help of a needle and then stained with carbol fuchsin for 15 min at room temperature (Solarbio). Coverslips were added, and the slides were observed with the light microscope (Olympus BX51, Olympus, Japan). Photographs were taken by the DP-73.

### Ultrastructure analysis by TEM

The microspore or pollen cell structure was studied by transmission electron microscopy (TEM). Anthers were fixed in 5% glutaraldehyde in phosphate-buffered saline (PBS) at 4 °C overnight, rinsed in 0.1 M PBS for three times, and post-fixed in 1% OsO_4_ (dissolved in 0.1 M PBS) for 3 h. After washing three times with PBS, the materials were then dehydrated with 50%, 70%, and 90% ethyl alcohol, 1/2 90% ethyl alcohol:1/2 acetone (V: V) and pure acetone for 15 min respectively. Anthers were embedded in Spurr’s resin for the cross-sectioning. Ultrathin sections (70 nm) were cut using a diamond knife on a Leica Ultracut ultramicrotome (Leica EM UC6). Sections were stained by saturated uranyl acetate and lead citrate and then examined with a transmission electron microscope (JEM1230, JEOL, Japan).

### Cytomembrane analysis by DiI staining

Pollen grains at stage 5 were collected in a 1.5 mL Eppendorf tube, hydrolyzed with enzyme solution containing 3% cellulose, 1% pectinase and 0.6% D-mannitol, and then by incubating at 35 °C for 3 h [36]. The hydrolyzed pollen grains were further stained with 10 μM 1’-dioctadecyl-3,3,3’,3’-tetramethylindocarbocyanine perchlorate (DiI, Sigma, USA) dissolved in acetate buffer (pH 5.5) for 10 min and then observed with the Olympus BX51 under an excitation wavelength of 549 nm [37].

### Nuclear analysis by DAPI

The integrity and stability of the nuclei were analyzed by 4’, 6-diamidino-2-phenylindole (DAPI) (Beyotime, Shanghai, China). Microspores or pollen grains were squeezed from the anther and stained with DAPI (10 μg mL^-1^) solution prepared in Tris-HCl (0.1 mol L^-1^ pH 9.0). The DAPI-stained microspores were observed with a fluorescence microscope (Olympus BX51, Olympus) and photographed with a DP-73 (Olympus, Japan) [38].

### Anther wall observation by paraffin sectioning

The anther wall was studied by paraffin sectioning. Anthers at different developmental stages were collected and fixed in FAA (5 mL of formaldehyde and 5 mL of glacial acetic acid in 90 mL of 70% (V/V) ethyl alcohol) at 4 °C overnight, and then dehydrated for 2 h in an ethyl alcohol gradient of 80%, 85%, 90%, and 95%, followed by dehydration in 100% ethyl alcohol twice, each time for 1 h. The anther was then hyalinized in 1/2 xylene and 1/2 ethyl alcohol overnight, saturated gradually with paraffin at 37 °C until saturation, and then immersed in pure paraffin overnight. The saturated materials were embedded, sectioned at 5 μm, dewaxed and then dyed with safranin-Fast Green or toluidine blue.

## Results

### Flower developmental stages

In convenience of research and description, the flower buds of the hybrids were classified into nine developmental stages according to the development of the male gametophyte, as shown in Fig. 1. The microsporocyte at stages 1 to stage 2 were undergoing the process of meiosis, and microspore release happened before stage 3 (Fig. 1), followed by the cell wall deposition and microspore mitosis at stage 4. The microspores developed further at stages 5 to 7 and matured into bicellular pollen grains at stages 8 and 9 (Fig. 1).

### Pollen viability and germination capacity difference between F1-S and F1-D

Mature pollen of F1-D was proved to be sterile by in vitro culture and seeds set assay [33]. FDA-PI staining was used to further explore the microspore/pollen viability at the process of development. Viable cells exhibited green fluorescence due to the effect of FDA, while DNA of dead cells showed red fluorescence of PI staining. The results revealed that microspores/pollen grains at all developmental stages of F1-S fertile were stained green by FDA, which was typically exampled as that in Fig. 2. A. Thus the pollen of F1-S fertile were viable. In F1-D sterile, microspores at stage 2 to stage 4 exhibited green fluorescence (Fig. 2. B, C), which weakened during the lateral stages 5 to 7 (Fig. 2. D, E), and was totally absent in mature pollen (Fig. 2. F). The above results showed that the microspore/pollen of F1-D sterile was gradually inactivated at the developmental process.

**Fig. 2.**
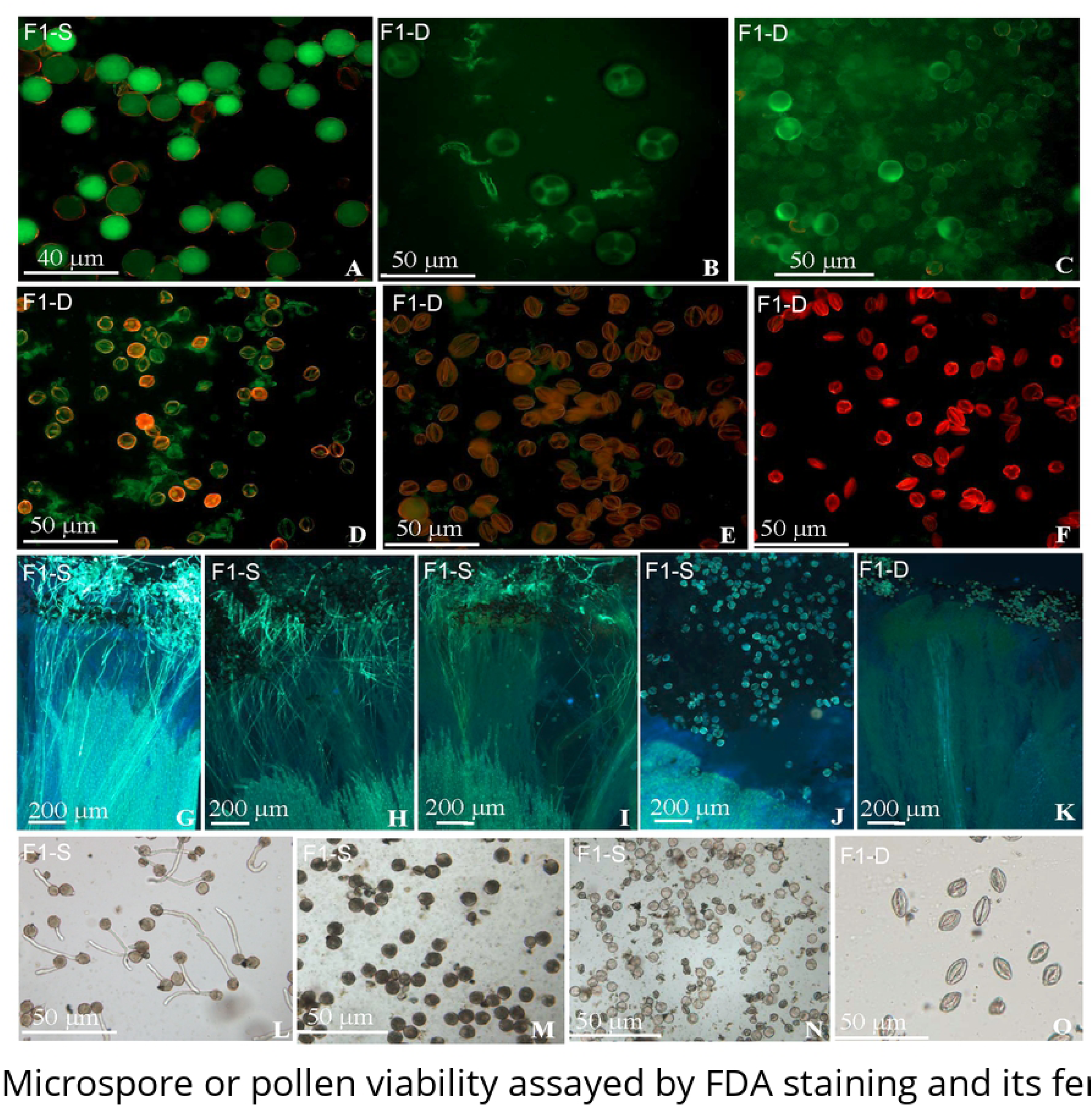
Microspore or pollen viability assayed by FDA staining and its fertility estimated by pollination and in vitro cultivation. A-F. Pollen viability assayed by FDA staining. A. Mature pollen grains of F1-S fertile; B. Microspore at stage 2; C. Pollen grains at stage 4; D. Pollen grains at stage 5; E. Pollen grains at stage 7; F. Pollen grains at stage 9; G-K. Pollen grains viability estimated by pollination; G-J show that the F1-S fertile pollen germination ratio was highest at stage 9; G. Pollen grains at stage 9; H. Pollen grains at stage 8; I. Pollen grains at stage 7; J. Pollen grains at stage 6; K. Pollen grains of F1-D sterile at stage 9; L. Pollen grains at stage 9 germinated well on in vitro medium; M, N. Pollen grains at stages 8 and 7 barely germinated on in vitro medium; O. Mature pollen grains (at stage 9) of F1-D sterile failed to germinate on in vitro medium.

Furthermore, microspore/pollen germination capacity was also analyzed at the developmental process (Fig. 1) by pollination and in vitro culture. In F1-S fertile, pollen grains at stages 7 to 9 germinated on the stigma (Fig. 2. G, H, I), and the germination ratio increased from stages 7 to 9, while pollen grains at stage 6 could not germinate on the stigma (Fig. 2. J). In F1-D sterile, microspores or pollen of all developmental stages failed to germinate on the stigma (Fig. 2. K). Pollen in vitro culture assay showed that pollen of F1-S fertile at stage 9 germinated well on the in vitro medium (Fig. 2. L), and the germination ratio was higher than 85%. However, the germination ratio was much lower for microspore at stages 7 and 8 (Fig. 2. M, N), and pollen at stage 6 could not germinate at all (Fig. 2. O). In F1-D sterile, pollen at all developmental stages was unable to germinate on in vitro medium.

The results of the FDA-PI staining, in vitro germination and pollination assays showed that microspores/pollen grains of F1-S fertile were viable at all developmental stages and fertile at maturation, which could be confirmed by seed set in F1-S fertile. While F1-D sterile pollen was gradually inactivated with development and incapable of germination. FDA-PI staining and the germination assay revealed that the pollen viability and germination capacity were not always consistent. Pollen with germination ability was inevitably positive for FDA-PI staining and viable, while viable pollen could only germinate at maturation.

### Abnormality in the process of microsporogenesis

Chromosome behavior during meiosis was vital for pollen fertility, and abnormal meiosis inevitably induces pollen abortion. Therefore, microsporogenesis at stages 1 to stage 2 was investigated by carbol fuchsin staining, to explore its causality with pollen abortion of F1-D sterile. The results showed that PMCs at stage 2 were undergoing meiosis I and meiosis II. Cytomixis (the migration of nuclear material from one PMC to another) was observed at zygotene of meiosis I in both F1-D sterile (Fig. 3. A, B) and F1-S fertile. A total of 120 PMCs at zygotene were analyzed for both F1-S fertile and F1-D sterile, and cells with cytomixis accounted for 7.0% and 18.57% of F1-S fertile and F1-D PMCs, respectively (Table 1). Furthermore, the chromosomes in the meiocytes of F1-D sterile were prone to leak at diakinesis of meiosis I (Table 1), implying that the chromosomes escaped out of the cytoplasm and the callose wall (Fig. 3. C, D). Based on an analysis of 140 PMCs at diakinesis in both the F1-S fertile and F1-D sterile, chromosome leakage accounted for 75.54% of the meiocytes of F1-D sterile, while only 2.00% meiocytes showed chromosome leakage in F1-S fertile (Table 1). Chromosome leakage even resulted in the occurrence of cytoplasts-cells that were completely devoid of nuclear material (Fig. 3. C, D). At the tetrad stage, abnormal and vestigial microspores were observed in the anther locule of F1-D sterile, and four microspores within the same callose wall were different in size and shape (Fig. 3. E), unlike those in F1-S fertile. Obviously, these results indicated that cytomixis and chromosome leakage occurred with higher frequency in the meiocytes of F1-D sterile.

**Table 1.** Chromosome and callose wall abnormalities during meiosis

|  | zygotene |  | diakinesis |  |
| --- | --- | --- | --- | --- |
|  | Meiocytes with asymmetrical callose wall (%) | Meiocytes with cytomixis (%) | Meiocytes with asymmetrical callose wall (%) | Meiocytes with chromosome extrusion (%) |
| F1-S-Fertile | 53.90±4.50 | 7.00±0.20 | 25.24±2.63 | 2.00±0.01 |
| F1-D-Sterile | 55.71±6.45 | 18.57±2.10 | 79.09±7.58 | 74.54±7.89 |

**Fig. 3.**
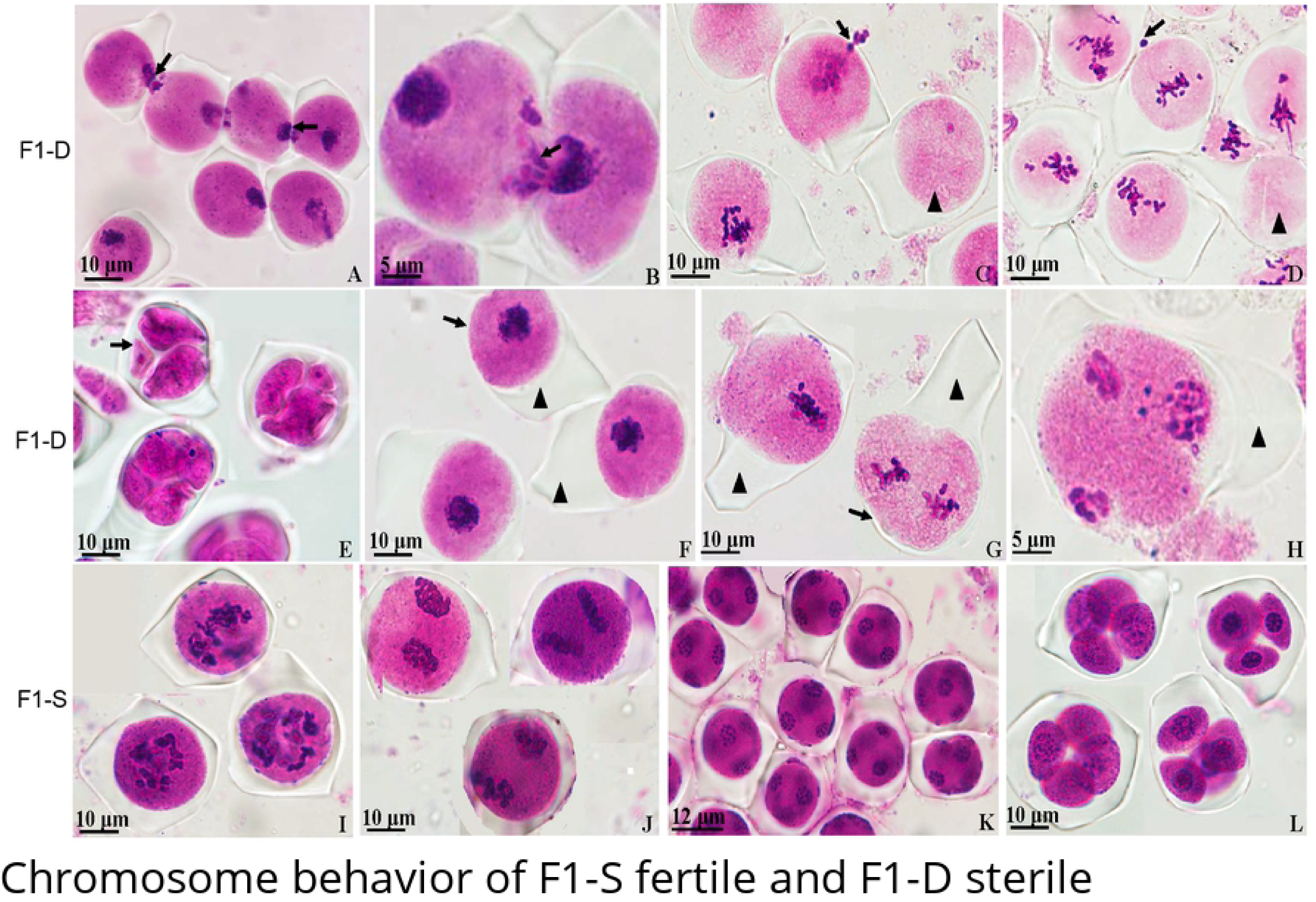
Chromosome behavior of F1-S fertile and F1-D sterile. A, B. Cytomixis at zygotene of meiosis I, with arrow showing nuclear material translocation between cells; C, D. Chromosome leakage at diakinesis of meiosis I, with arrow showing a chromosome escaping a meiocyte and arrowhead showing a PMC without chromosome; E. Asymmetrical callose wall deposition at the tetrad stage, with arrow indicating microspores of unequal size and morphology; F, G, H. Callose wall deposited asymmetrically at different developmental stages, with arrow and arrowhead showing thinner and thickened callose deposition spots around the meiocytes, respectively; I, J, K, and L show the callose wall deposited symmetrically at diakinesis and metaphase of meiosis I, at telophase of meiosis II, and at the tetrad stage.

Regular callose accumulation and dissolution are vital for male fertility, and the callose wall deposition in F1-S fertile and F1-D sterile was accordingly studied. The results revealed that asymmetrical callose wall deposition, resulting in thinned or perforated areas of the callose wall, was observed in both F1-S fertile and F1-D sterile (Fig. 3. F, G)) at zygotene of meiosis I, and occurred with high frequency in both types of meiocytes at the rates of 53.90% and 55.71%, respectively. However, at diakinesis, the percentage of asymmetrical callose wall deposition dropped to 25.24% in F1-S fertile, while increased to 79.09% in F1-D sterile (Table 1). In F1-D sterile, asymmetrical callose wall deposition occurred at a high percentage at all developmental stages before microspore release (Fig. 3. A-H), while the callose wall was deposited relatively symmetrically after diakinesis in F1-S fertile (Fig. 3. I, J, K, L). Asymmetrical callose deposition was also observed via TEM in the PMCs of F1-D sterile (Fig. 5. C). These results indicated that asymmetrical callose wall deposition occurred with higher frequency in F1-D sterile. Nuclear material translocation and leakage occurred in the areas of thinned or perforated callose wall deposition in the meiocytes (Fig. 3. A, B, C).

**Fig. 4.**
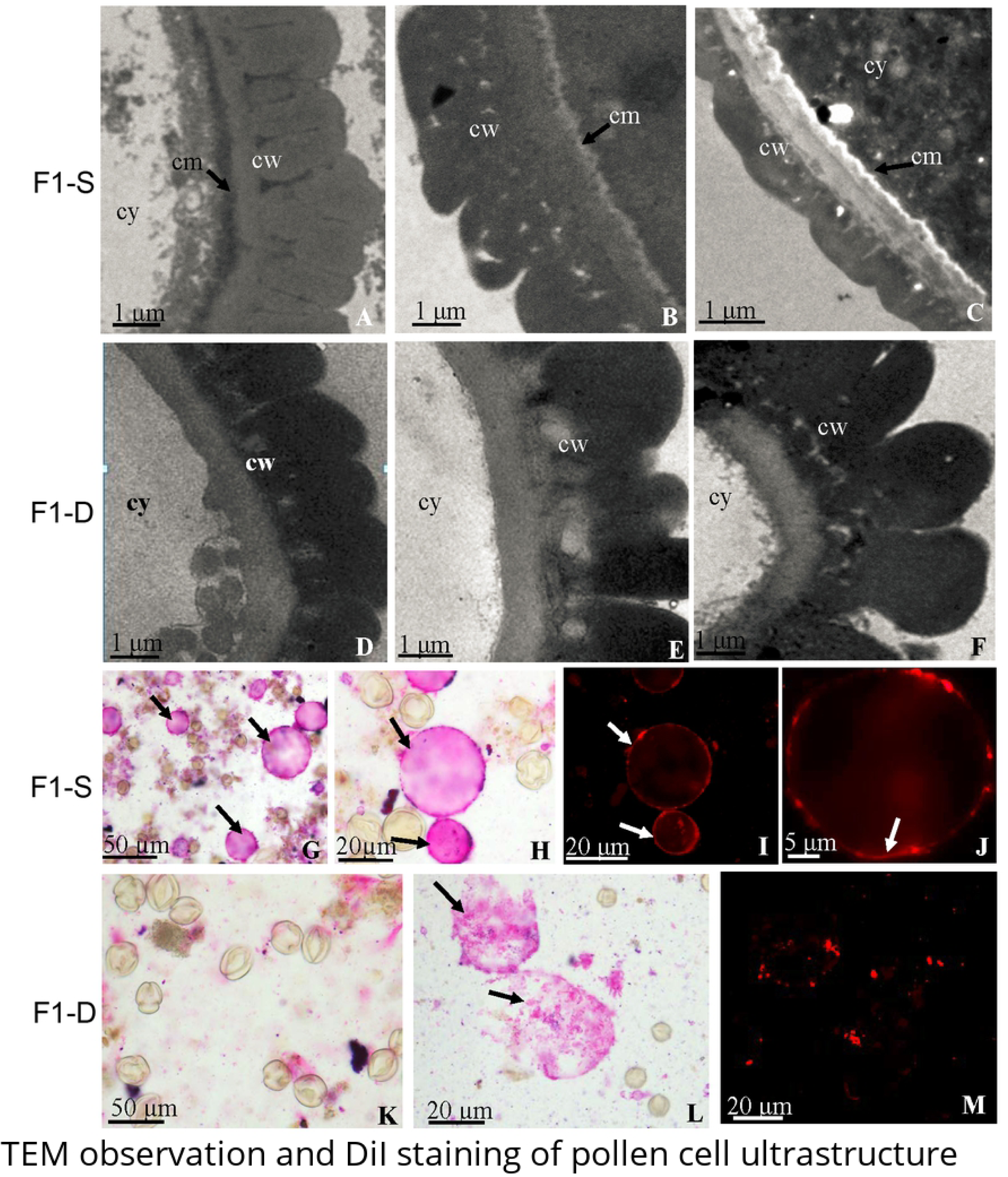
TEM observation and DiI staining of pollen cell ultrastructure. A, B, C. Detailed cell wall structure of F1-S fertile, with arrow indicating the cytomembrane; D, E, F. Detailed cell wall structure of F1-D sterile pollen deficient in cytomembrane; A, D. Pollen cell wall structure at stage 3; B, E. Pollen cell wall structure at stage 5; C, F. Pollen cell wall structure at stage 9; G-M. Observation of cytomembrane by DiI staining; G, H. Mature pollen of F1-S fertile hydrolyzed with cellulase and pectinase, with arrow showing the protoplast; I, J. Mature pollen of F1-S fertile stained with DiI, with arrow showing the cytomembrane; K, L. Mature pollen of F1-D sterile hydrolyzed with cellulase and pectinase, with arrow showing the ruptured protoplast; M. Sterile pollen negative for DiI staining. al, anther locule; cm, cell membrane; cy, cytoplasm; cw, cell wall.

**Fig. 5.**
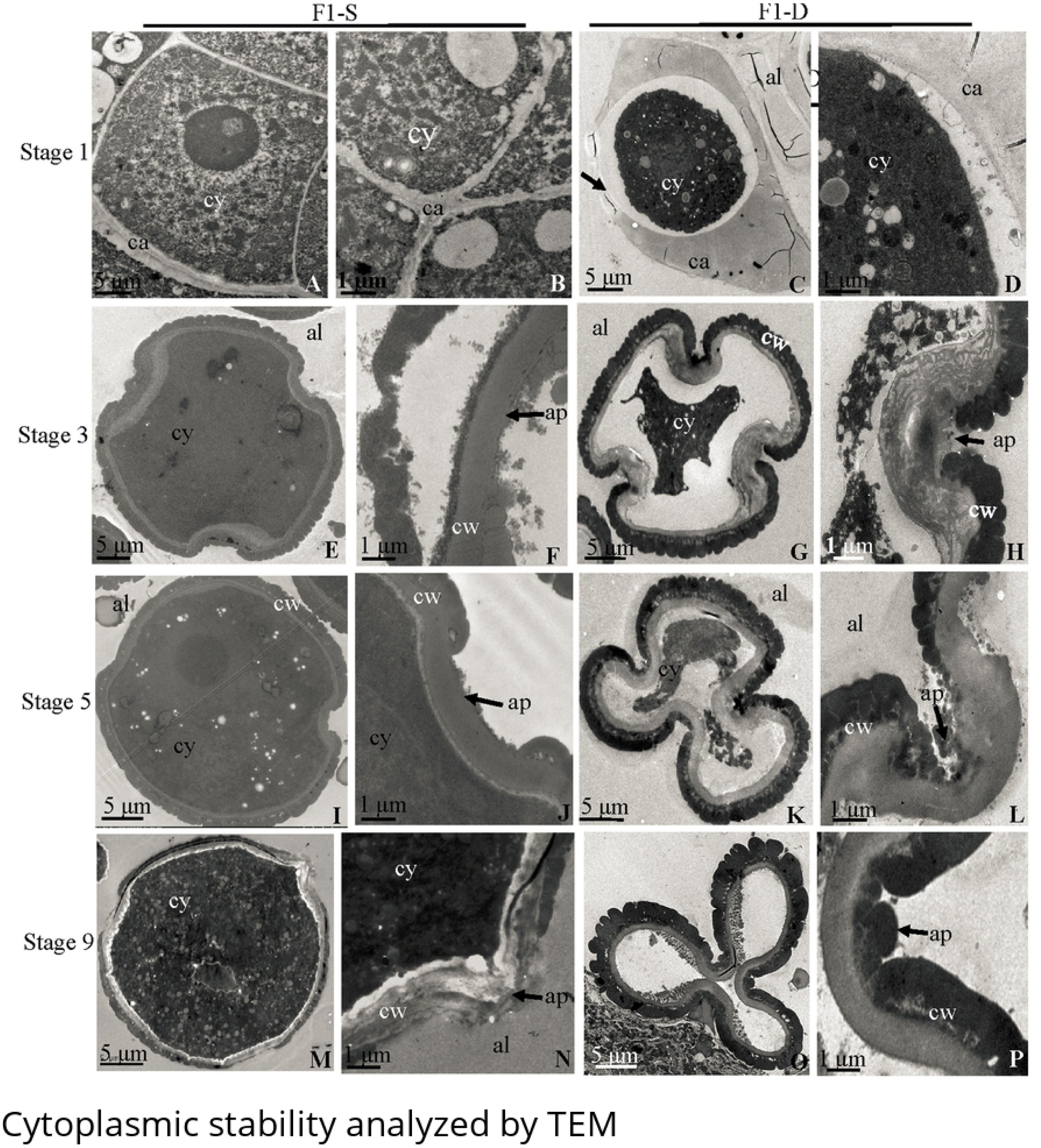
Cytoplasmic stability analyzed by TEM. A. The PMC was full of cytoplasm; B. The cytoplasm of PMC was tightly covered by the callose layer; C. The arrow indicated the perforated area resulted from asymmetric callose wall deposition; D. The gap between the cytoplasm and callose wall of PMC; E. The microspore was full of cytoplasm; F. The arrow indicated that there was no sporopollenin deposition in the aperture region; G. The cytoplasm was partially extruded; H. The arrow showed the sporopollenin was not deposited in the aperture region; I. The microspore was full of cytoplasm; J. The arrow indicated that sporopollenin did not deposited in the aperture region; K. The pollen gradually collapsed in the aperture region; L. The arrow showed that the sporopollenin was gradually deposited in the aperture region; M. The pollen was full of cytoplasm; N. The arrow showed that there was no sporopollenin deposition in the aperture region; O. The pollen grain was totally absent of cytoplasm, and collapsed at the aperture region; P. The arrow showed that the aperture region was totally blocked by sporopollenin. ap, aperture; al, anther locule; ca, callose; cm, cell membrane; cy, cytoplasm; cw, cell wall.

### Cytomembrane deficiency and pollen wall deposition abnormality in microspore/pollen of F1-D

Abnormality in the cell membrane might interfere with the movement of substances in and out of cells and with the development of the pollen wall structure [24], and aberrant pollen wall formation might also lead to pollen abortion. Microspore/pollen was further studied with TEM, to reveal the cell structure differences between F1-S fertile and F1-D sterile. The most significant difference was that the cell membrane was well developed in the microspores or pollen of F1-S fertile at all represented developmental stages, and became electron-translucent from stage 3 to maturity (Fig. 4. A, B, C); however, the cytomembrane was undetectable in F1-D sterile, as shown in microspores at stage 3, 5 and 9 (Fig. 4. D, E, F). The cell membrane deficiency was further validated by DiI staining, which revealed that the protoplasts of the F1-D sterile pollen grains ruptured immediately after hydrolysis, and the cytomembrane was indiscernible after DiI staining (Fig. 4. K, L, M), which might reduce the stability of protoplast and be responsible for the above pollen rupture. In contrast, intact and rounded protoplasts of various sizes were hydrolyzed from F1-S fertile pollen grains, and the cytomembrane was positively dyed red (Fig. 4. G, H, I, J).

The cell wall of fertile and sterile pollen was both composed of the exine (nexine and sexine) and intine. In F1-D sterile pollen, the exine did not develop in the aperture region at stage 3 (Fig. 5. G, H), and sporopollenin gradually deposited in this region at stage 5 (Fig. 5. K, L). The aperture of the mature pollen was totally covered by the exine (Fig. 5. O, P). In fertile pollen, only intine was deposited in the aperture region at the above three developmental stage (Fig. 5. F, J, N).

### Nuclear and cytoplasmic instability in pollen of F1-D

As cytomembrane deficiency was observed in the microspore/pollen of F1-D sterile, the cell inclusion stability was further studied. TEM observation revealed that the cytoplasm of PMCs in F1-S fertile were covered tightly by the callose cell wall (Fig. 5. A, B), while there was gap between the cytoplasm and callose wall of PMCs in F1-D sterile (Fig. 5. C, D). PMCs of F1-D sterile had abundant cell inclusions (Fig. 5. C), which were extruded gradually at later developmental stages (Fig. 5. G, K), and the cytoplasm was totally absent from mature pollen grains (Fig. 5. O). The pollen of F1-D sterile also collapsed gradually around the apertures, resulting in three apertures close to each other at the cell center of the mature pollen (Fig. 5. O). In F1-S fertile, the cytoplasm, organelles and amyloid became richer from stage 1 to the mature pollen stage, and the mature pollen was plump, regular, and filled with abundant cell inclusions (Fig. 5. A, E, I, M).

DAPI staining was used to further examine the microspore/pollen nuclei stability at stage 2 to stage 9 (Fig. 1), which indicated that micronuclei were present in the tetrad phase and the following monokaryotic phase in F1-D sterile (Fig. 6. A, B). The released microspores underwent one round of mitosis and generated binucleate pollen grains at stage 4 (Fig. 6. C). In F1-D sterile, two nuclei were present in the microspore at stage 4, absent from some pollen grains at stage 6 (Fig. 6. D), and absent from all pollen grains at stages 7 to 9 (Fig. 6. E, F). In F1-S fertile, two nuclei also arose at stage 4, but they were always present at all the subsequent stages; therefore, mature F1-S fertile pollen had two nuclei (Fig. 6. G), while none was present in F1-D sterile pollen. DAPI analysis further provided evidence that the microspore/pollen nuclei of F1-D sterile were not stable during development.

**Fig. 6.**
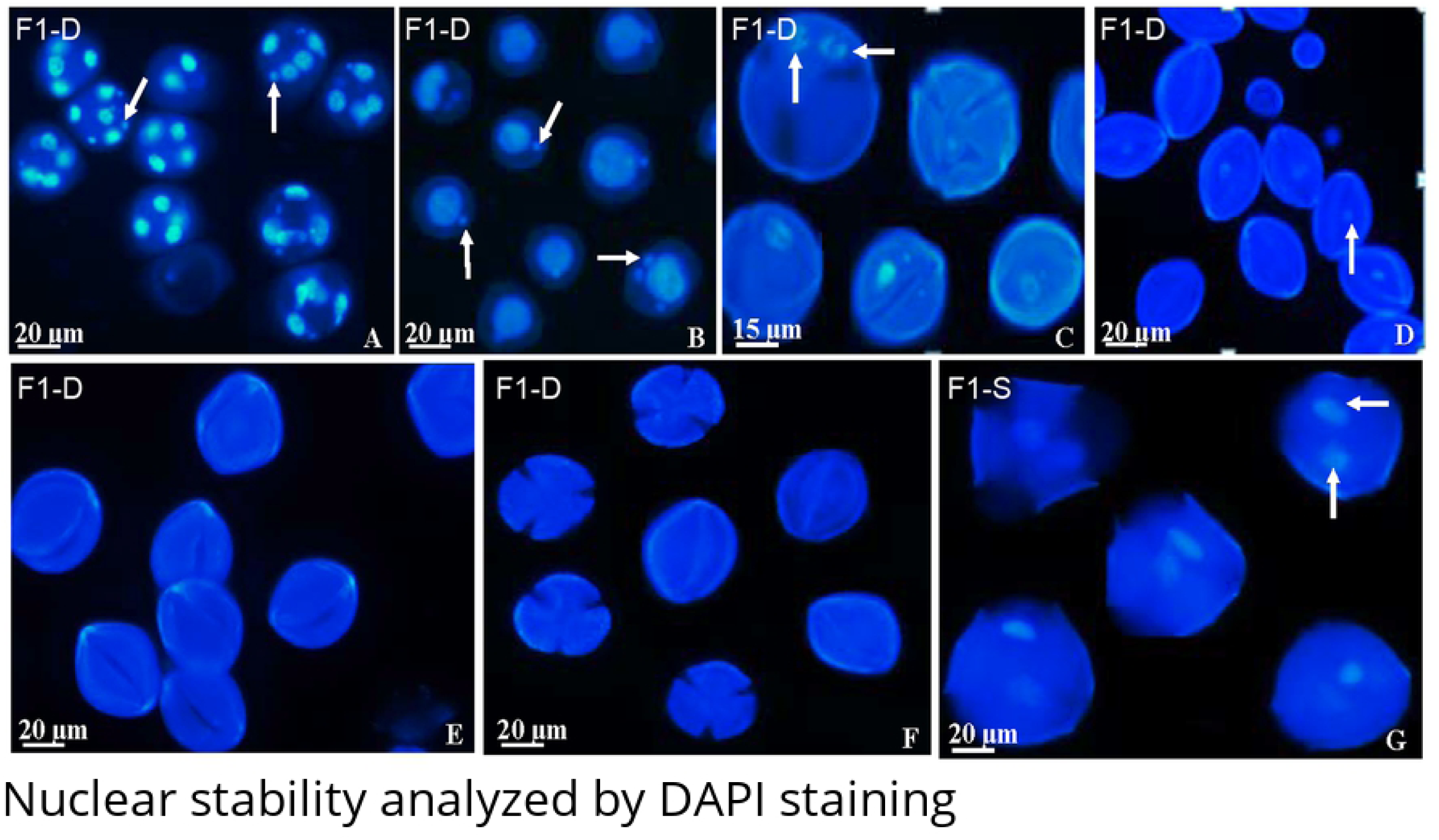
Nuclear stability analyzed by DAPI staining. A, B. Micronucleus at the tetrad phase and the following monokaryotic phase, with arrow indicating the micronucleus; C. The arrow indicated two nuclei at stage 4; D. Nuclei disappeared in some pollen grains at stage 6; E, F. Nuclei disappeared in all pollen grains at stages 7 and 9; G. Two nuclei in the mature pollen of F1-S fertile.

### Abnormal tapetum development in F1-D

The tapetum plays major roles in pollen wall development and pollen viability. Observation of the anther wall structures in the hybrids showed that the anthers developed normally in F1-S fertile, with normal connectivum and four anther locules of equal size (Fig. 7. A). In F1-D sterile, about 66% of the anthers were normal, as described above (Fig. 7. B), while others had enlarged connectivum and reduced anther locules (Fig. 7. C, D), which might morphologically affect microsporogenesis and pollen development. The anther wall was composed of epidermis, fibrous endothecium, one middle layer and the tapetum, which formed normally in both the sterile and fertile hybrids (Fig. 7. E, F) at the beginning of PMC differentiation. For F1-S fertile hybrids, the tapetum started to degenerate at the early tetrad stage (Fig. 7. G) and mostly disappeared at the release of the microspores (Fig. 7. H). However, the tapetum in F1-D sterile degenerated later than that in F1-S fertile. The tapetum was intact at the tetrad stage (Fig. 7. I), mostly remained after the release of the free microspores (Fig. 7. J, K), and ultimately degenerated at stage 5 (Fig. 7. L). Therefore, the tapetum degradation was delayed in the anther of F1-D sterile.

**Fig. 7.**
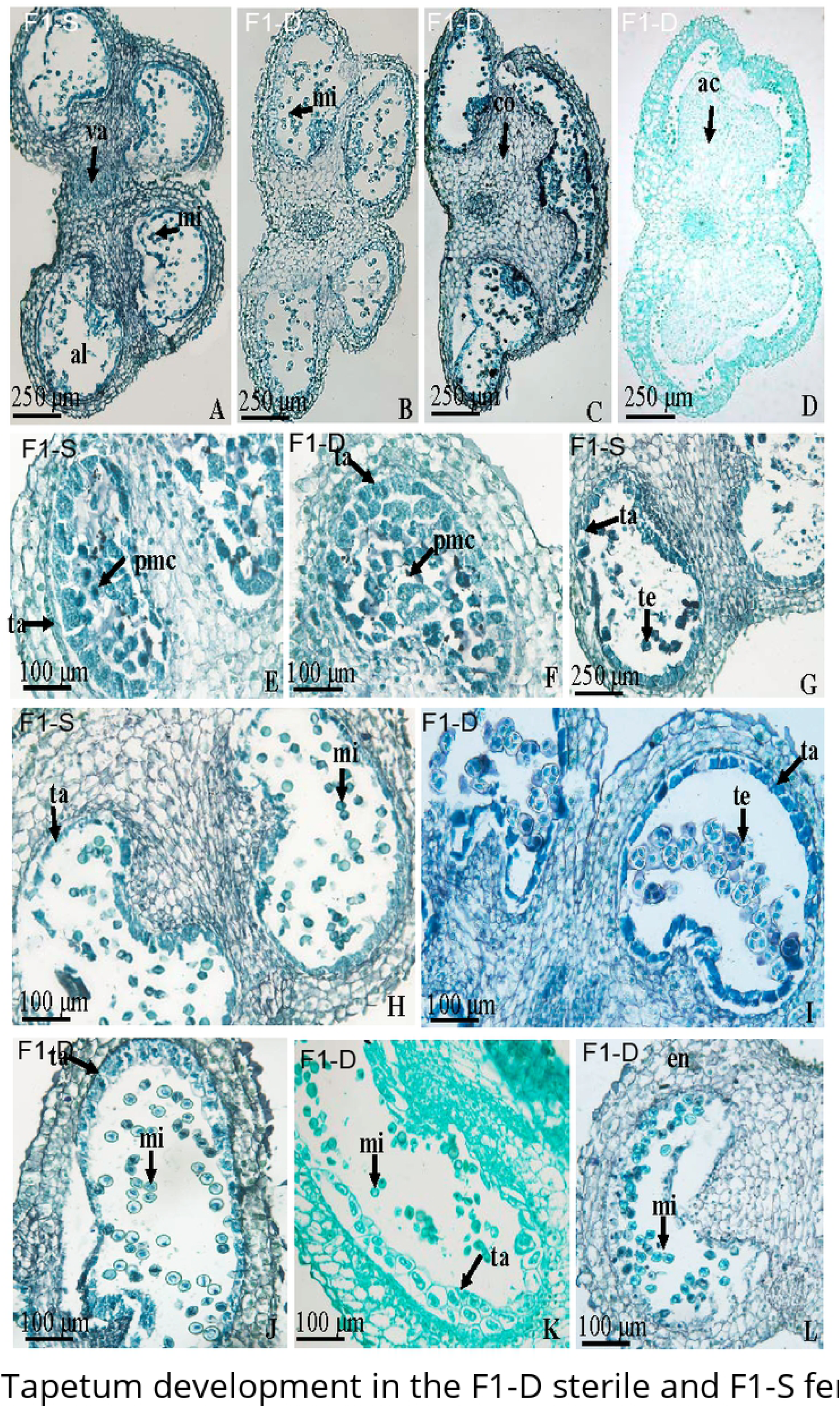
Tapetum development in the F1-D sterile and F1-S fertile hybrids. A. Anther of F1-S fertile at stage 3; B, C, D. Anther of F1-D sterile; B. Anther with normal connectivum and four locules at stage 3; C. Anther with enlarged connectivum and three locules at stage 4; D. Anther with inflated connectivum and only two locules at stage 4; E. Tapetum formation in F1-S fertile; F. Tapetum formation in F1-D sterile; G. The tapetum of F1-S fertile began to degrade at the tetrad stage; H. The tapetum of F1-S fertile was almost entirely degraded by the release of the microspores; I. The tapetum of F1-D sterile was almost intact at the tetrad stage; J, K. The tapetum of F1-D sterile partially presented at the release of the microspore; L. The tapetum of F1-D sterile was degraded at stage 5. co, connectivum; en, endothecium; mi, microspore; pmc, pollen mother cell; ta, tapetum; te, tetrad; va, vascular bundle.

## Discussion

Sterility in interspecific hybrids constrains their application in breeding improvement and has long been an interesting biological topic [39, 40]. Microsporogenesis and microspore development are under strict control by the surrounding sporophytic tissue and the gametophytic intracellular environment. In this study, we investigated the cytological basis of pollen sterility of hybrid obtained from a cross between a *Nicotiana* CMS maternal parent and *N. alata* [33]. The results showed that cytomixis, chromosome leakage, irregular callose wall deposition occurred with higher frequency in the male gametophyte of F1-D sterile hybrid, the cytomembrane was deficient, and the tapetum degradation was delayed.

Cytomixis has been reported to occur along cytomictic channels [26, 27, 41, 42], and cytomixis during the prophase of the first meiotic division is widespread in plant species and is evolutionarily important [43-46]. Our study showed that cytomixis occurred at zygotene of meiosis I in both the PMCs of F1-D sterile and that of F1-S fertile. However, the cytomixis ratio in the PMCs of F1-D sterile was 11.57% higher than that in the PMCs of F1-S fertile. It is reported that irregular cytomixis may be caused by various disturbances such as the structure of the cytomictic channels, synthesis of the callose coat, cell wall formation, the structure of the cell cytoskeleton and genetic unbalance [27, 42]. As has been reviewed, the microtubular apparatus does not play an obvious role in the process of cytomixis in tobacco [26, 42, 47, 48]. Furthermore, asymmetrical deposition of the callose wall occurred at similar ratio at the zygotene of meiosis I in the PMCs of both the F1-S fertile and F1-D sterile hybrids. Therefore, the high percentage of asymmetric callose wall deposition at the beginning of meiosis might not result in higher cytomixis in F1-D sterile, but only provide convenience or a prerequisite for the transfer of nuclear material between PMCs during meiosis. Genetic unbalance might be partially responsible for higher cytomixis in F1-D sterile. Furthermore, cytomembrane deficency observed in this reseach might also be related with higher cytomixis in the PMCs of F1-D sterile.

The asymmetrical callose wall deposition percentage declined at diakinesis of meiosis I in F1-S fertile, while it increased in F1-D sterile, revealing that callose wall rearrangement happened during the process of meiosis. This result was consistent with the report that callose wall deposition occurred almost continuously throughout meiosis, ceasing only with the final compartmentalization of microspores after telophase II [24]. The results also showed that chromosome leakage occurred 75% more frequently in F1-D sterile than that in F1-S fertile and was always linked with thinned or perforated areas of the callose wall. The high percentage of chromosome leakage in the sterile pollen might be due to a cell membrane deficiency. Moreover, it has also been reported that callose wall deposition at the microspore surface is mediated by the plasma membrane [49]. We deduced that this cell membrane deficiency might be responsible for the asymmetrical callose wall deposition at diakinesis in the PMCs of F1-D sterile, which then provided a convenient path for chromosome leakage. Chromosome leakage inevitably intensified genetic material distribution abnormalities during meiosis, as reflected by the presence of deformed microspores and micronuclei at the tetrad stage and the monocyte stage.

In this study, microspore and pollen instability in the F1-D sterile was also found. DAPI staining and TEM analysis showed that the cytoplasm and nucleus were extruded from the pollen grain during the processes of microspore maturation. This phenomenon could also be attributed to the plasma membrane deficiency and genetic unbalance of the sterile pollen. In addition, the results also indicated that sporopollenin was irregularly deposited in the aperture region of F1-D sterile pollen, which might be induced by delayed tapetum degeneration after microspore release.

The timely degradation of the tapetum is necessary for pollen development and fertility, and the timing of tapetum degeneration varies among species. For example, degeneration commenced at the tetrad stage and was completed at the bicellular pollen stage in *Brachypodium* and rice [11, 50]. However, in wheat, barley and *Actinidia deliciosa*, the breakdown of tapetum cells began during the vacuolated microspore stage [11, 34, 51]. In this study, tapetum degradation in F1-D sterile was delayed compared with that in F1-S fertile, which might partially cause the observed pollen incapacity.

The genetic basis of hybrid sterility has been studied in some plants, including *Solanum*, rice (*Oryza*), *Mimulus, Arabidopsis lyrata*, and *A. thaliana*, revealing that male sterility in these species was either induced by large-scale chromosome organization changes (including chromosome rearrangement) or merely by genic loci (including two loci with Bateson–Dobzhansky–Muller interactions and one locus representing a male sterility factor) [36-38, 52]. In this research, pollen sterility in F1-D sterile might be both genetically and morphologically regulated. Higher cytomixis and chromosome leakage might be related with the genetic unbalance, and the cytomembrane deficiency might also be responsible for the instability of the nuclear and cytoplasm material. Furthermore, abnormal tapetum degradation might also result in the pollen abortion in F1-D sterile. The cytomembrane deficiency observed in F1-D sterile might be the result of lower expression level of the putative nuclear fertility restoration genes [33], which worked insufficiently in restore pollen fertility in F1-D hybrids.

## Acknowledgements

Not applicable

## Reference

1. Zhang ZS, Lu YG, Liu XD, Feng JH, Zhang GQ. Cytological mechanism of pollen abortion resulting from allelic interaction of F-1 pollen sterility locus in rice (*Oryza sativa* L.). Genetica. 2006; 127(1-3): 295–302. doi: 10.1007/s10709-005-4848-z. PubMed PMID: WOS: 000239164900026.

2. Long Y, Zhao L, Niu B, Su J, Wu H, Chen Y, et al. Hybrid male sterility in rice controlled by interaction between divergent alleles of two adjacent genes. Proceedings of the National Academy of Sciences of the United States of America. 2008; 105(48): 18871–6. doi: 10.1073/pnas.0810108105. PubMed PMID: WOS: 000261489100048.

3. Nikova V, Vladova R, Pundeva R, Euphytica DSJ. Cytoplasmic male sterility in *Nicotiana tabacum* L. obtained through interspecific hybridization. Euphytica. 1997; 94(3): 375–8. doi: 10.1023/a:1002941527164.

4. Yang YX, Li YH, Tong JF, Qasim SM, Chen ZX, Wang L et al. Wide-compatibility gene S5n exploited by functional molecular markers and its effect on fertility of intersubspecific rice hybrids. Crop Science. 2012; 52(2): 669–75. doi:10.2135/cropsci2011.04.0232

5. Zhao ZG, Jiang L, Zhang WW, Yu CY, Zhu SS, Xie K et al. Fine mapping of S31, a gene responsible for hybrid embryo-sac abortion in rice (*Oryza sativa* L.). Planta. 2007; 226: 1087–96. doi: 10.1007/s00425-007-0553-8. PubMed PMID: 17549514

6. Goldberg RB, Beals TP, Sanders PM. Anther development: Basic principles and practical applications. Plant Cell. 1993; 5:1217–29 doi: 10.1105/tpc.5.10.1217. PMID: 8281038

7. Scott RJ, Speilman M, Dickinson HG. Stamen structure and function. Plant Cell. 2004, 16: S46–S60. doi: 10.1105/tpc.017012. PMID: 15131249

8. Garcia CC. Anther wall formation in Solanaceae species. Annals of botany. 2002; 90(6): 701–6. doi: 10.1093/aob/mcf248. PubMed PMID: MEDLINE:12451025.

9. Gómez JF, Talle B, Wilson ZA. Anther and pollen development: A conserved developmental pathway. Journal of Integrative Plant Biology. 2015; 57: 876–91. doi: 10.1111/jipb.12425 PMID: 26310290

10. Pacini E, Franchi GG, Hesse M. The tapetum: Its form, function, and possible phylogeny in Embryophyta. Plant Systematics & Evolution. 1985;149(3-4):155-85. doi: 10.1007/bf00983304.

11. Xu J, Ding Z, Vizcay-Barrena G. Aborted microspores acts as a master regulator of pollen wall formation in *Arabidopsis*. Plant Cell. 2014; 26(4):1544–56. doi: 10.1105/tpc.114.122986. PubMed PMID: WOS: 000337351300014.

12. Sarkar S, Bhattacharyya S, Gantait S. Cytological analysis for meiotic patterns in wild rice (*Oryza rufipogon* Griff.). Biotechnology reports (Amsterdam, Netherlands). 2017; 13: 26–9. doi: 10.1016/j.btre.2016.12.004. PubMed PMID: MEDLINE: 28352559.

13. Simeonova E, Wypiórkiewicz E, Charzyńska M. Pollen development in *Cucumis sativus* L. Acta biologica Cracoviensia. Series botanica. 1999; 41: 139–42.

14. Akanksha S, Singh MB, Bhalla PL. Ultrastructure of microsporogenesis and microgametogenesis in *Brachypodium distachyon*. Protoplasma. 2015; 252: 1575–86. doi: 10.1007/s00709-015-0793-6. PMID: 25772681

15. Lu P, Chai M, Yang J, Ning G, Wang G, Ma H. The *Arabidopsis* callpse defective microspore1 gene is required for male fertility through regulating callose metabolism during microsporogenesis. Plant Physiology. 2014;164(4):1893–904. doi: 10.1104/pp.113.233387. PubMed PMID: WOS: 000334342800030.

16. Waterkeyn, L. Les parois microsporocytaires de nature callosique chez Helleborus et Tradescantia. Cellule. 1962; 62: 225–55.

17. Waterkeyn L, Bienfait A. On a possible function of the callosic special wall in *Ipomoea purpurea* (L) Roth. Grana. 1970; 10: 13–20. doi: 10.1080/00173137009429852

18. Dong XY, Hong ZL, Sivaramakrishnan M, Mahfouz M, Verma DPS. Callose synthase (CalS5) is required for exine formation during microgametogenesis and for pollen viability in *Arabidopsis*. Plant Journal. 2005; 42(3): 315–28. doi: 10.1111/j.1365-313X.2005.02379.x. PubMed PMID: WOS: 000228488500003.

19. Sun XH, Zhang ZG, Feng D, Zhang Q, Han LD, Wu JX et al. Glucan synthase-like 5 (GSL5) plays an essential role in male fertility by regulating callose metabolism during microsporogenesis in rice. Plant and Cell Physiology. 2015; 56(3): 497–509. doi: 10.1093/pcp/pcu193. PubMed PMID: WOS: 000352494100010.

20. Leofanti GA, Camadro EL. Pollen viability and meiotic abnormalities in brome grasses (*Bromus* L., Section Ceratochloa) from Argentina. Turkish Journal of Botany. 2017; 41(2):127–33. doi: 10.3906/bot-1607-46. PubMed PMID: WOS: 000398038400002.

21. Mwathi MW, Gupta M, Atri C, Banga SS, Batley J, Mason AS. Segregation for fertility and meiotic stability in novel *Brassica allohexaploids*. Theoretical and Applied Genetics. 2017; 130(4):767–76. doi: 10.1007/s00122-016-2850-8. PubMed PMID: WOS: 000398737400011.

22. Haque SM, Ghosh B. Cell division study in ‘pollen mother cells and pollen grains of in vivo and ex vitro plants in *Drimiopsis botryoides*. Grana. 2017; 56(2): 124–36. doi: 10.1080/00173134.2016.1144785. PubMed PMID: WOS: 000394651300003.

23. Soares TL, de Souza EH, Costa, MAPD, Silva, SDE, dos Santos-Serejo, JA. Viability of pollen grains of tetraploid banana. Bragantia. 2016; 75(2): 145–51. doi: 10.1590/1678-4499.328. PubMed PMID: WOS: 000378221000003.

24. Larson DA, Skvarla JJ. Fine structural studies of *Zea Mays* pollen: cell membranes and exine ootogeny. American Journal of Botany. 1996; 53: 1112–25. doi:10.1002/j.1537-2197.1966.tb06879.x

25. Junqueira VB, Costa AC, Boff T, Mueller C, Correa Mendonca MA et al. Pollen viability, physiology, and production of maize plants exposed to pyraclostrobin plus epoxiconazole. Pesticide Biochemistry and Physiology. 2017; 137: 42–8. doi: 10.1016/j.pestbp.2016.09.007. PubMed PMID: WOS: 000399067000007.

26. Mursalimov S, Sidorchuk Y, Deineko E. Behavior of nucleolus in the tobacco male meiocytes involved in cytomixis. Cell Biology International. 2017; 41(3): 340–4. doi: 10.1002/cbin.10718. PubMed PMID: WOS: 000394887700012.

27. Xie B, Deng Y, Kanaoka MM, Okada K, Hong Z. Expression of *Arabidopsis* callose synthase 5 results in callose accumulation and cell wall permeability alteration. Plant Science. 2012; 183: 1–8. doi: 10.1016/j.plantsci.2011.10.015. PubMed PMID: WOS: 000299713300001.

28. Zhao B, Shi H, Wang W, Liu X, Gao H, Wang X et al. Secretory CopII protein SEC31B is required for pollen wall development. Plant Physiology. 2016; 172(3): 1625–42. doi: 10.1104/pp.16.00967. PubMed PMID: WOS: 000391172300021.

29. Liu YG, Chen L. Male sterility and fertility restoration in crops. Annu Review Plant Biology. 2014; 65: 579–606. doi: 10.1146/annurev-arplant-050213-040119. PMID: 24313845.

30. Miedaner T, Herter CP, Gosslau H, Wilde P, Hackauf B. Correlated effects of exotic pollen-fertility restorer genes on agronomic and quality traits of hybrid rye. Plant Breeding. 2017; 136(2): 224–9. doi: 10.1111/pbr.12456. PubMed PMID: WOS: 000398638700013.

31. Bharaj TS, Virmani SS, Khush GS. Chromosomal location of fertility restoring genes for ‘wild abortive’ cytoplasmic male sterility using primary trisomics in rice. Euphytica. 1995; 83: 169–73. doi:10.1007/BF01678126.

32. Mallick EH. Induced cytoplasmic male sterility and fertility restoration and production of hybrid rice. Genetica Agraria. 1980; 207–13.

33. Liao JG, Dai JR, Yang SY, Zhou XL, Ren LH, Chen ZZ et al. Interspecific cross-hybrids of *Nicotiana tabacum* L. cv. (*gla*.) S ‘K326’ with *Nicotiana alata*. Plant Breeding. 2017; 136(3): 427–35. doi: 10.1111/pbr.12474. PubMed PMID: WOS: 000403463700017.

34. Giuseppina F, Simone DA, Rita B. Tapetum and middle layer control male fertility in *Actinidia deliciosa*. Annals of Botany. 2013; 112: 1045–55. doi: 10.1093/aob/mct173. PMID: 23965617

35. Gómez JF Wilson ZA. A barley PHD finger transcription factor that confers male sterility by affecting tapetal development. Plant Biotechnology Journal. 2014; 12: 765–77. doi: 10.1111/pbi.12181. PMID: 24684666

36. Moyle LC, Graham EB. Genetics of hybrid incompatibility between *Lycopersicon esculentum* and *L. hirsutum*. Genetics. 2005;169(1): 355–73. doi: 10.1534/genetics.104.029546. PubMed PMID: WOS: 000227039900030.

37. Moyle LC, Nakazato T. Comparative genetics of hybrid incompatibility: sterility in two Solanum species crosses. Genetics. 2008; 179(3): 1437–53. doi: 10.1534/genetics.107.083618. PubMed PMID: WOS: 000258313400025.

38. Simon M, Durand S, Pluta N, Gobron N, Botran L, Ricou A et al. Genomic conflicts that cause pollen mortality and raise reproductive barriers in *Arabidopsis thaliana*. Genetics. 2016; 203: 1353–67. doi: 10.1534/genetics.115.183707. PubMed PMID: WOS: 000379473600025

39. Maheshwari S, Barbash DA. The Genetics of Hybrid Incompatibilities. Annual Review of Genetics. 2010; 45(1): 331–55. doi: 10.1146/annurev-genet-110410-132514.

40. Novo, PE, Valls, JFM, Galdeano F. Interspecific hybrids between *Paspalum plicatulum* and *P. oteroi*: a key tool for forage breeding. Crop Science. 2016; 73: 356–62. doi: 10.2135/cropsci2010.10.0610.

41. Kravets EA. Cytomixis and its role in the regulation of plant fertility. Russian Journal of Developmental Biology. 2013; 44: 113–28. doi: 10.1134/s1062360413030028. PubMed PMID: WOS: 000318928700001.

42. Sidorchuk IuV, Deineko EV, Shumnyi VK. The role of microtubular cytoskeleton and callose walls in the cytomixis process in tobacco (*Nicotiana tabacum* L.) pollen mother cells. Tsitologiia. 2007; 49: 876-80. PubMed PMID: MEDLINE: 18074779.

43. Kamra OP. Chromatin Extrusion and Cytomixis in Pollen Mother Cells of Hordeum. Hereditas. 1960; 46: 592–600. doi: 10.1111/j.1601-5223.1960.tb03103.x

44. Basavaiah, Murthy TCS. Cytomixis in pollen mother cells of *Urochloa panicoides* P. Beauv. (*Poaceae*). Cytologia. 1987; 52: 69–74. doi:10.1508/cytologia. 52. 69

45. FeijÓ JA, Pais MSS. Cytomixis in meiosis during the microsporogenesis of *Ophris lutea*: an ultrastructural study. Caryologia. 1995; 42: 37–48.

46. Caetano C, Pagliarini MS. Cytomixis in maize microsporocytes. Cytologia. 1997; 62: 351–55. doi:10.1508/cytologia.62.4_351.

47. Zadoo SN, Choubey RN, Gupta SK, Premachandran, MN. Chromosomal Stability in the Backcross Progenies of Pentaploid Hybrids between *Avena sativa* L. and *A. maroccana* Gdgr. Plant Breeding. 2010; 100: 316–9. doi:10.1111/j.1439-0523.1988.tb00258.x.

48. Pierozzi NI, Moura MF. Cytological analyses in ‘Niagara Branca’ grape and in its somatic mutant ‘Niagara Rosada’. Notulae Botanicae Horti Agrobota. 2014; 42: 460–5. doi:10.15835/nbha.42.2.9540.

49. Blackmore S, Wortley AH, Skvarla JJ, Rowley JR. Pollen wall development in flowering plants. New Phytologist. 2007; 174: 483–98. doi:10.1111/j.1469-8137.2007.02060.x PMID: 17447905

50. Onyemaobi I, Liu H, Siddique KHM, Yan G. Both male and female malfunction contributes to yield reduction under water stress during meiosis in bread wheat. Frontiers in Plant Science. 2017; 7. doi: 10.3389/fpls.2016.02071. PubMed PMID: WOS: 000391503700002.

51. Gómez JF Wilson ZA. A barley PHD finger transcription factor that confers male sterility by affecting tapetal development. Plant Biotechnology Journal. 2014; 12: 765–77. doi: 10.1111/pbi.12181 PMID: 24684666

52. He JH, Shahid MQ, Li YJ, Guo HB, Cheng XA, Liu XD, et al. Allelic interaction of F-1 pollen sterility loci and abnormal chromosome behaviour caused pollen sterility in intersubspecific autotetraploid rice hybrids. Journal of Experimental Botany. 2011; 62(13): 4433–45. doi: 10.1093/jxb/err098. PubMed PMID: WOS: 000294968400004.

